# Implication of synaptotagmins 4 and 7 in activity-dependent somatodendritic dopamine release

**DOI:** 10.1101/2021.01.25.427983

**Authors:** Benoît Delignat-Lavaud, Charles Ducrot, Willemieke Kouwenhoven, Nina Feller, Louis-Éric Trudeau

**Affiliations:** Department of Pharmacology and Physiology, Faculty of Medicine, Université de Montréal; Department of Neurosciences, Faculty of Medicine, Université de Montréal; CNS Research Group (GRSNC), Montréal, QC, Canada H3C 3J7

## Abstract

Dopamine (DA) neurons can release DA not just from axon terminals, but also from their somatodendritic (STD) compartment thought a mechanism that is still incompletely understood. Using voltammetry in mouse mesencephalic brain slices, we find that STD DA release has low capacity, is stable in response to electrical but not optogenetic train pulses and shows a calcium sensitivity that is comparable to that of axonal release. It is also strikingly more resilient compared to axonal release in a 6‐ hydroxydopamine model of Parkinson’s disease plasticity. We find that the molecular mechanism of STD DA release differs from axonal release with regards to the implication of synaptotagmin (Syt) calcium sensors. While individual constitutive knock-out Syt4 and Syt7 is not sufficient to reduce STD DA release, removal of both isoforms reduces this release by ~50%, leaving axonal release unimpaired. Our works unveils clear differences in the mechanisms of STD and axonal DA release.

## INTRODUCTION

Dopamine (DA) neurons of the mesencephalon play a key role in motor control, motivated behaviors and cognition^1,2^. DA neurons can release DA not only from axon terminals by a classical exocytosis mechanism^3^, but also through their somatodendritic (STD) compartment, as demonstrated by multiple approaches including *in vivo* microdialysis, fast scan cyclic voltammetry (FSCV) and patch-clamp recordings of D2 receptor mediated currents in the ventral tegmental area (VTA) and substantia nigra pars compacta (SNc)^4–7^. These nuclei contain the cell body and dendrites of DA neurons, but little if any DA-containing axon terminals^5,8^. Although there is limited direct evidence, STD DA release is believed to be implicated in regulating the excitability of DA neurons though activation of STD D2 autoreceptors^9^. It has also been suggested to regulate motor behaviors^10,11^, mainly by local activation of D1 receptors.

The molecular mechanism of STD DA release is still unclear. Reversal of the DA transporter (DAT) has been proposed^12^, but this mechanism cannot account for the results of studies that measured STD DA release *in vitro* and *in vivo* in the presence of DAT blockers. These studies unequivocally show that blocking DAT leads to an increase in extracellular DA, whether in evoked release^4,6,9,13,14^ or spontaneous release^15–20^. A vesicular exocytotic-like mechanism has therefore been proposed, in agreement with the fact that STD DA release is activity-dependent (TTX-sensitive)^9,18,21^, reserpine-sensitive^9,22,23^, calcium-dependent^9,13,18,20,22,24^, and blocked by botulinum neurotoxins, which disrupt SNARE-proteins^18,25,26^. Although large pools of DA-containing small clear synaptic vesicles are not found in the dendrites of DA neurons, these dendrites contain pleiomorphic vesicles that bear the vesicular monoamine transporter (VMAT2), suggesting that they could be sites of DA storage in dendrites^27^. Together, these findings suggest that, although there may be some fundamental differences between the mechanisms of terminal and STD DA release, both implicate a form of exocytosis.

Although STD DA release is calcium-dependent, conflicting results exist regarding the calcium-sensitivity of STD DA release in comparison with axonal release. Previous studies performed in guinea pig reported that STD DA release persists at extracellular calcium concentrations as low as 0.5 mM, a concentration at which axonal release is typically abrogated from most axon terminals^13,18^. In contrast, previous work performed with mouse tissue and indirectly detecting STD DA release using the patch-clamp technique and STD D2 receptor activation, reported that axonal and STD DA release display a similar calcium-dependency^9,24,28–31^. Here, we reexamined this question in mouse brain slices after optimizing direct detection of DA using FSCV.

STD DA release could play a role in adaptations of basal ganglia circuitry and motor behaviors during the progression of Parkinson’s disease (PD). Compatible with the dying-back hypothesis of PD suggesting that PD pathology starts at the axon terminal level^32^, previous work in rats measuring baseline STD DA release by microdialysis suggested that this form of release, is preserved for longer periods of time compared to axonal DA release following 6-OHDA lesions^33^. It is unclear if activity-dependent STD release is similarly resilient. Here, we examined the impact of a striatal 6-hydroxydopamine (6-OHDA) lesion on evoked STD DA release 1 day after the lesion, where an early loss of axon terminals occurs, and 14 days after the lesion at a time where soma and dendrites of DA neurons are severely impacted^34^.

Finally, an important outstanding question is the identification of the molecular mechanisms of STD DA release. Prior work has demonstrated that many proteins involved in regulated exocytosis, such as the calcium sensor Syt1, are selectively targeted to the axonal domain of neurons and not in dendrites^3,19,35–37^. Building on previous *in vitro* work suggesting possible roles of Syt4 and Syt7^19^, in the present study we tested the hypothesis that Syt4 and Syt7 play a key role in STD DA release in the intact brain by quantifying STD DA release in Syt4, Syt7 and Syt4/7 double knockout (KO) mice.

## MATERIALS AND METHODS

### Animals

Male and female mice of 11-12 weeks were used in this study. For optogenetic experiments, B6;129S-Gt(ROSA)26Sor^tm32(CAG‑COP4*H134R/EYFP)Hze^/J (Ai32, The Jackson Laboratory, stock 012569, USA) homozygote mice expressing a floxed H134R variant of the light-activated channelrhodopsin-2 were bred with homozygote B6.SJL-Slc6a3^tm1.1(cre)Bkmn^/J (DAT^IREScre^, The Jackson Laboratory, stock 006660, USA) expressing the cre recombinase under control of the DAT promoter, allowing channelrhodospin-2 to be expressed selectively in DA neurons. Heterozygote DAT^IREScre^ mice were also used for experiments in which ChR2 was virally expressed. Constitutive knock-out mice for Syt4 (129S6.129X1(B6)-Syt4^tm1Hahe^/J, The Jackson Laboratory, stock #012400, USA)^38^, Syt7^39^ and WT littermates were bred from heterozygous crosses or crossed with each other to obtain double KO mice. Genotyping for Syt4 KO mice was determined using specific primers to target the wild type Syt4 sequence (primers Syt4WT-fwd and Syt4WT-rev) and the neomycin cassette within the mutated allele (primers neo-fwd and Syt4WT-rev) – Syt4WT-fwd: CACTTCCCTCACGTCAGAGGAG, - Syt4WT-rev: GCAAGGAGAGCTCTTGGATGTG, - neo-fwd: AACCACACTGCTCGACATTGGG. Genotyping for Syt7 KO mice was performed using specific primers to target the wild type Syt7 sequence (Syt7WT-fwd: CATCCTCCACTGGCCATGAATG; - Syt7WT-rev: GCTTCACCTTGGTCTCCAG) and the neomycin cassette within the mutated allele (neo-fwd: CTTGGGTGGAGAGGCTATTC; neo-rev: AGGTGAGATGACAGGAGATC), as provided by Jackson. Genotyping for Syt7 mutation in combined Syt4/7 KO mice was determined using another set of specific primers due to overlapping sequences within the neomycin cassette used in both the Syt4 and Syt7 mouse lines: - neo-fwd: CTTGGGTGGAGAGGCTATTC and Syt7WTexon4: AGTGTCCAGGCTCCC. Experiments were performed blind with regards to animal genotype, with the exception of Syt4 KO mice, because these KO mice could be easily identified due to a neurodevelopmental alteration of the anterior commissure and corpus callosum (**Fig. S1**). All procedures involving animals and their care were conducted in accordance with the Guide to care and use of Experimental Animals of the Canadian Council on Animal Care. The experimental protocols were approved by the animal ethics committees of the Université de Montréal. Housing was at a constant temperature (21°C) and humidity (60%), under a fixed 12h light/dark cycle, with food and water available ad libitum.

### Stereotaxic injections

6-7 week-old DAT^IREScre^ mice were anesthetized with isoflurane (Aerrane; Baxter, Deerfield, IL, USA) and fixed on a stereotaxic frame (Stoelting, Wood Dale, IL, USA). A small hole was drilled in the exposed skull and a Hamilton syringe was used for the injections. For optogenetic experiments, an adeno-associated virus (AAV5-EF1a-DIO-hChR2(H134R)-EYFP, 4,2×10^12^ vg/mL, UNC GTC Vector Core, USA) was injected bilaterally at the following injection coordinates [AP (anterior–posterior; ML (medial– lateral); DV (dorsal-ventral), from bregma], to infect neurons in the entire ventral mesencephalon: AP −3.0 mm; ML +/− 1.0 mm; DV −4.5 mm. Animals recovered in their home cage and were closely monitored for 3 days. The animals were used one month after injection, allowing maximal expression of ChR2 in DA neurons. Success of the injection was visually validated each time during the slicing of the brains by visualizing the presence of the eYFP reporter. For 6-OHDA experiments, saline or 6-OHDA (5 μg/μL; 2 μL in total, at a rate of 0.5 μL/min, Sigma, Canada) were injected unilaterally in the dorsal striatum: AP +1.0 mm; ML +1.5 mm; DV −2.8 mm. The brains were used for FSCV experiments 1 or 14 days after the injection.

### Brain slice preparation and solutions

Acute brain slices from 11-12-week-old male or female mice were used for the FSCV recordings. When possible, matched pairs of WT and KO mice were used on each experimental day. The animals were anesthetized with halothane, quickly decapitated and the brain harvested. Next, the brain was submersed in ice-cold oxygenated artificial cerebrospinal fluid (aCSF) containing (in mM): NaCl (125), KCl (2.5), KH_2_PO_4_ (0.3), NaHCO_3_ (26), glucose (10), CaCl_2_ (2.4), MgSO_4_ (1.3) and coronal VTA and/or striatal brain slices of 300 μm thickness were prepared with a VT1000S vibrating blade microtome. Once sliced, the tissue was transferred to oxygenated aCSF at room temperature and allowed to recover for at least 1h. For recordings, slices were placed in a custom-made recording chamber superfused with aCSF at 1 ml/min and maintained at 32°C with a TC-324B single channel heater controller (Warner Instruments, USA). All solutions were adjusted at pH 7.35-7.4, 300 mOsm/kg and saturated with 95% O_2_-5% CO_2_ at least 30 min prior to each experiment.

### Fast scan cyclic voltammetry recordings

Optically or electrically evoked DA release was measured by FSCV using a 7 μm diameter carbon-fiber electrode placed into the tissue ~100 μm below the surface. A bipolar electrode (Plastics One, Roanoke, VA, USA) or an optical fiber connected to a 470 nm wavelength LED was placed ~200 μm away. Carbon-fiber electrodes were fabricated as previously described^40^. Briefly, carbon fibers (Goodfellow Cambridge Limited, UK) of 7 μm in diameter were aspirated into ethanol-cleaned glass capillaries (1.2 mm O.D., 0.68 mm I.D., 4 inches long; World Precision Instruments, FL, USA). The glass capillaries were then pulled using a P-2000 micropipette puller (Sutter Instruments, Novato, USA), dipped into 90°C epoxy for 30s (Epo-Tek 301, Epoxy Technology, MASS, USA) and cleaned in hot acetone for 3s. The electrodes were heated at 100°C for 12h and 150°C for 5 days. Electrodes were polished and filed with potassium acetate at 4M and potassium chloride at 150 mM. The protruding carbon fibers were cut using a scalpel blade under direct visualization to a length allowing to obtain maximal basal currents of 100 to 180 nA.

The electrodes were calibrated with 1 μM DA in aCSF before and after each recorded slice and the mean of the current values obtained were used to determine the amount of released DA. After use, electrodes were cleaned with isopropyl alcohol (Bioshop, Canada). The potential of the carbon fiber electrode was scanned at a rate of 300 V/s according to a 10 ms triangular voltage wave (−400 to 1000 mV vs Ag/AgCl) with a 100 ms sampling interval, using a CV 203BU headstage preamplifier (Molecular Devices) and a Axopatch 200B amplifier (Molecular Devices, USA). Data were acquired using a Digidata 1440a analog to digital converter board (Molecular Devices, USA) connected to a computer using Clampex (Molecular Devices, USA). Slices were left to stabilize for 20 min before any electrochemical recordings. After positioning of the bipolar stimulation electrode or the optical probe and carbon fiber electrodes in the tissue, single pulses (400 μA or 30 mW, 1ms,) or pulses-train (30 pulses at 10 Hz) were applied to the tissue to trigger DA release. For evaluating the calcium dependency of axonal and STD release, variations of calcium concentrations in the aCSF (0, 0.5 and 2.4 mM) were compensated by changing the concentration of MgSO_4_ to keep divalent cation levels equivalent.

### Immunohistochemistry

For Syt immunolabelling experiments, 40 μm brain slices from animals perfused with 4% paraformaldehyde (in PBS, pH-7.4) were cut with a cryostat (Leica CM 1800; Leica Canada) and used for immunohistochemistry (IHC). Because selective and specific Syt4, Syt7 and VMAT2 antibodies were all from the same host species (rabbit), a double labeling protocol (Jackson ImmunoReseach) with monovalent Fab fragments was used. After a PBS wash, the tissue was permeabilized, nonspecific binding sites blocked (goat serum 5%) and incubated overnight at room temperature with the first primary antibody (rabbit anti-Syt1, anti-Syt4 or anti Syt7 from Synaptic Systems, Germany; 1:1000), followed by 2h with a first secondary antibody (rabbit Alexa Fluor-488–conjugated, 1:500, Invitrogen, Canada). A blocking step of antigenic sites from the first primary and secondary antibody combination was performed thereafter by a 3h incubation with normal serum from the same species as the primary antibody, followed by a blocking solution (goat block: PBS, Triton X100 0.3%, bovine serum albumin 5%) with 50 μg/mL of unconjugated monovalent Fab fragments against the host of the primary antibody, overnight, at room temperature and under agitation. Slices were then washed, and a second labeling was performed with a second primary antibody (rabbit anti-VMAT2, 1:1000, gift of Dr. Gary Miller, Colombia University), and a second secondary antibody (rabbit Alexa Fluor-546–conjugated, 1:500, Invitrogen). For each IHC staining, a control group was included, with the full protocol except for omission of the second primary antibody. A classical immunostaining protocol was used for the knockout validation of Syt4 and Syt7 antibodies (**Fig. S2**), using mouse anti-tyrosine hydroxylase (Millipore Sigma; 1:1000) and rabbit anti-Syt4 or anti-Syt7 primary antibodies (Synaptic Systems; 1:1000) subsequently detected using Alexa Fluor-488-conjugated and Alexa Fluor-546-conjugated secondary antibodies (Invitrogen; 1:500).

### Confocal Imaging

Images were acquired using an Olympus Fluoview FV1000 point-scanning confocal microscope (Olympus, Canada) with a 60x oil-immersion objective (NA 1.35). Images acquired using 488nm and 546 nm laser excitation were scanned sequentially to prevent non-specific bleed-through signal. All image analysis was performed using ImageJ (National Institutes of Health) software.

### Reverse Transcriptase-quantitative PCR

We used RT-qPCR to quantify the amount of mRNA encoding Syt1, 4, 5, 7 and 11 in brain tissue from P70 Syt4^+/+^ and Syt4^-/-^ mice and P70 Syt7^+/+^ and Syt7^-/-^ mice. Adult whole brains were harvested and homogenized in Trizol solution, then RNA extraction was performed using RNAeasy Mini Kit (Quiagen, Canada) according to the manufacturer’s instructions. The concentration and purity of the RNA from DA neurons were determined using a NanoDrop 1000 (Thermo Scientific, Waltham, MA USA). Total purified RNA (40 ng) was reverse-transcribed in a total of 20 μl including 1 μl of dNTP, 1 μl of random hexamer, 4 μl of 5X buffer 5X, 2 μl of dithiothreitol (DTT), 1 μl of RNAse-Out and 1 μl of the Moloney Murine Leukemia Virus reverse transcriptase enzyme (MML-V, Invitrogen). Quantitative PCR was carried out in a total of 15μl consisting of 3μl cDNA, 7.5μl SYBER green PCR master mix (Quanta Biosciences, USA), 10μM of each primer, completed up to 15μl with RNA-free water. qPCR was performed on a Light Cycler 96 machine (Roche, Canada) using the following procedure: 10 min at 95°C; 40 cycles of 30s at 95°C, 40s at 57°C and 40s at 72°C; 1 cycle of 15s at 95°C, 15s at 55°c and 15s at 95°C. Results were analysed with Light Cycler 96 software and Excel. The efficiency of the reaction (E=10^(−1/slope)^ −1) was calculated from the slope of the linear relationship between the log values of the RNA quantity and the cycle number (Ct) in a standard curve. Calculation of relative mRNA levels was performed by using the 2^(−DDCt) formula^41^, where the Ct value of the mRNA level for Syt1, 4, 5, 7 and 11 was normalized to the Ct value of GAPDH in the same sample. Ct values used were the mean of duplicate repeats. Melt-curves of tissue homogenate indicated specific products after Syt1, 4, 5, 7 and 11 qPCR mRNA amplification, attesting of the adequate quality of the primers chosen (not shown). Primers were designed with the Primers 3 and Vector NTI software and were synthetized by Alpha DNA (Montreal, QC). Primers for qPCR were as follows: Syt1: 5’ GTGGCAAGACACTGGTGAT 3’ and 5’ CTCAGGACTCTGGAGATCG 3’; Syt4: 5’ CACTTCCCTCACGTCAGAGGAG 3’ and 5’ GCAAGGAGAGCTCTTGGATGTG 3’; Syt5: 5’ GTCCCATACGTGCAACTAGG 3’ and 5’ AACGGAGAGAGAAGCAGATG 3’; Syt7: 5’ CCAGACGCCACACGA 3’ and 5’ CCTTCCAGAAGGTCT 3’; Syt11: 5’ CTTGTATGGCGGGGTCTTGT 3’ and 5’ ATACGCCCCAGCTTTGATGA 3’ and GAPDH: 5’ GGAGGAAACCTGCCAAGTATGA 3’ and 5’ TGAAGTCGCAGGAGACAACC 3’. Primers were tested by comparing primers sequences to the nucleotide sequence database in GenBank using BLAST (www.ncbi.nlm.nih.gov/BLAST/).

### Statistics

Data are presented as mean +/− SEM. The level of statistical significance was established at p < 0.05 in one-way ANOVAs with appropriate post-hoc tests and two-tailed t tests, performed with Prism 8 software (GraphPad, *p < 0.05, **p < 0.01, ***p < 0.001, # p < 0.0001).

## RESULTS

### D2 autoreceptors and DAT limit the extent of somatodendritic dopamine release in mouse VTA slices

The difficulty to reliably detect STD DA release in mouse rodent slices has greatly slowed progress in better understanding the mechanisms and roles of this form of DA release. We therefore first aimed to optimize its detection in mouse VTA slices by comparing different modes of stimulation and physiological parameters that may limit its extent.

Previous studies performed in brain slices or *in vivo* typically triggered STD DA release using extracellular electrical stimulation^42–44^. A downside of this approach is that it non-selectively depolarizes local afferent terminals and the cell bodies of local GABA and glutamate neurons in addition to DA neurons. In recent years, optical stimulation using channelrhodopsin-2 (ChR2) or other opsin variants has increasingly been used to obtain more selective activation of DA neuron axons^45^. However, to this date, this approach has not been used to selectively trigger STD DA release in FSCV experiments. We first evaluated whether optogenetic stimulation of DA neurons might be more effective to trigger STD DA release or produce more stable release (**Fig. 1A, 1B**).

**Figure 1:**
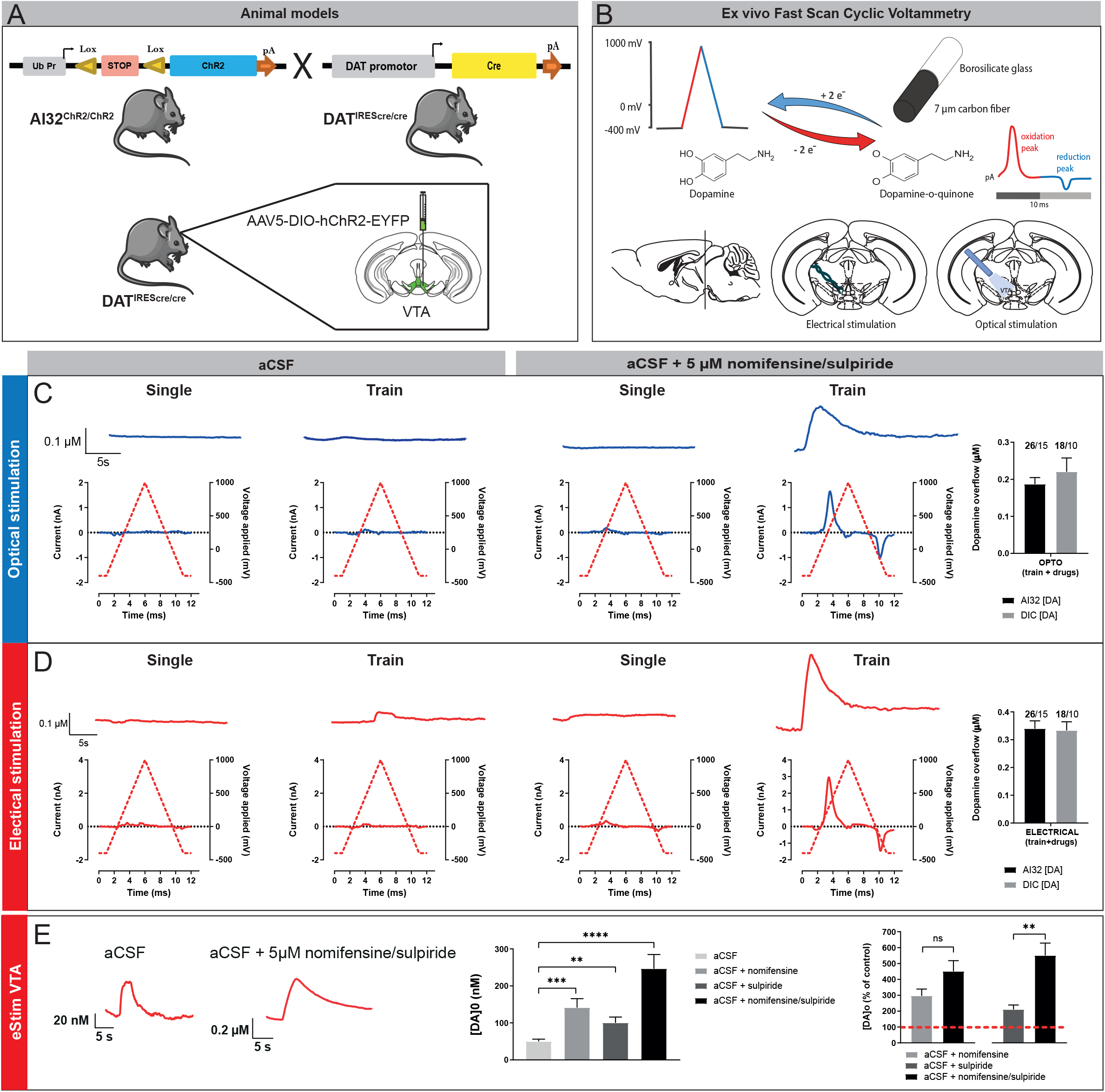
Optogenetic and electrical stimulation trigger comparable levels of somatodendritic dopamine release in mouse VTA slices. **(A)** Animal models used for optogenetic experiments. We either used a mouse line expressing a floxed version of light-activated channelrhodopsin (ChR2) crossed with a DA-specific Cre driver line (DAT^IREScre^) or performed stereotaxic injections of AAV5-EF1a-DIO-hChR2(H134R)-eYFP virus in the VTA of DAT^IREScre^ mice to selectively express ChR2 in DA neurons. **(B)** Fast Scan Cyclic Voltammetry was used to monitor DA levels. A voltage ramp of −400 to 1000 mV vs Ag/AgCl at 300 V/s was used, with a 100 ms sampling interval. Recordings were made in coronal slices containing the VTA and DA release was triggered by either optical stimulation with a 470 nm blue light LED or with a bipolar stimulating electrode. **(C)** Top, representative traces of responses obtained in the VTA with 1 pulse (1ms) of blue light («single pulse») or a pulse-train of stimulation (30 pulses of 1 ms at 10 Hz), in the presence of normal ACSF or ACSF + a DAT blocker (nomifensine, 5 μM) and an antagonist of D2 autoreceptors (sulpiride, 5 μM). Bottom, voltammograms of the representative traces. **(D)** Top, representative traces of responses obtained in the dorsal striatum with 1 electrical pulse (1 ms, 400 μA) or a pulse-train (30 electrical pulses of 1 ms at 10 Hz, 400 μA), in the presence of ACSF or ACSF + 5 μM nomifensine/sulpiride. Bottom, voltammograms of the representative traces. **(E)** Effect of nomifensine/sulpiride on STD DA release measured by pulse-train electrical stimulation.

We compared single pulse optical (1 ms, 470 nm) and electrical (1 ms, 400 μA) stimulation. Recordings were performed in the VTA of DAT^IREScre^/Ai32 mice, in which ChR2 is conditionally expressed in all DA neurons, and in DAT^IREScre^ heterozygote mice injected in the VTA with a floxed hChR2-EYFP AAV construct. Neither stimulation conditions, either in normal ACSF or in the presence of DAT (nomifensine, 5 μM) and D2 receptor blockade (sulpiride, 5 μM) yielded detectable evoked elevations of extracellular DA (**Fig. 1C, 1D**). However, the use of pulse trains (30 pulses at 10 Hz) in the presence of nomifensine and sulpiride allowed reliable detection of STD DA release in VTA slices, both for electrical stimulation (average peak DA levels of 340 nM +/− 28 nM in DAT^IREScre^/Ai32 mice [n = 15] and 333 nM +/− 32 nM in DAT^IREScre^ mice infected with ChR2 AAV [n = 10]) and for optical stimulation (average peak DA levels at 187 nM +/− 17 nM in DAT^IREScre^/Ai32 mice [n = 15] and 220 nM +/− 37 nM in DAT^IREScre^ mice infected with ChR2 AAV [n = 10]). Although peak levels of activity-dependent STD DA release in the two strains of mice tended to be higher with electrical compared to optical stimulation, the difference between the two modes of stimulation was not significantly different (**Fig. 1C, 1D**, right bar graphs).

Reuptake through the DAT and the D2 autoreceptor are two well-known regulators of extracellular DA levels and DA release^46–48^. We next examined the effect of DAT and D2 receptor blockade individually to determine whether a combined block of reuptake and autoreceptor function was required to reliably detect STD DA release in response to electrical train stimulation. Each recording was performed after 15 min of nomifensine or sulpiride or a combination of the two. Baseline levels of evoked DA in the absence of antagonist were very small, but still reliably detectable (49 nM +/− 6 nM). Blockade of DAT or D2 receptors individually, caused a significant increase in the maximal amplitude of evoked STD DA release in the VTA (+296% +/− 42% for nomifensine alone [n = 6], +210% +/− 28% for sulpiride alone [n = 6]), while a combination of the two drugs caused a cumulative increase of 500% +/− 51% [n = 12], thus demonstrating that the two manipulations were mostly additive and that a combined blockade of both membrane proteins allowed to maximally increase the detected signal.

### Optogenetic stimulation reveals strong use-dependent attenuation of evoked STD DA release in the VTA

In previous work evaluating STD DA release using FSCV in guinea pig brain slices, repeated stimuli were found to cause stimulation-dependent attenuation^5^. This represents a major limiting factor to further examine the mechanisms of this form of release. Using optical train stimulation in the presence of nomifensine and sulpiride, we therefore evaluated the stability of STD DA release in response to a series of 7 consecutive stimuli with an interstimulus interval of 5 min. STD DA overflow evoked by optical stimulation showed a robust and progressive decrease in peak amplitude in response to repeated stimuli (**Fig. 2A**). By the end of the stimulation protocol, evoked STD DA release decreased by approximately 50% in DAT^IREScre^/Ai32 mice and by approximately 40% in virally transduced DAT^IREScre^ mice. Compatible with the possibility that this decrement was due to rundown of releasable pools of DA, a 20 min delay before a final stimulation revealed a clear partial recovery. Strikingly, electrical stimulation failed to cause a similarly extensive rundown of STD DA, with only a modest, non-significant decrease of less than 20% detected by the last of 7 stimuli (**Fig. 2B**). Although speculative, this lack of rundown in response to electrical stimulation could be due to the recruitment of afferent fibers that secrete neuromodulators able to maintain vesicular DA stores for longer periods of time. Due to this favorable characteristic of electrical stimulation on the stability of STD DA release, all further experiments were performed with this mode of stimulation.

**Figure 2:**
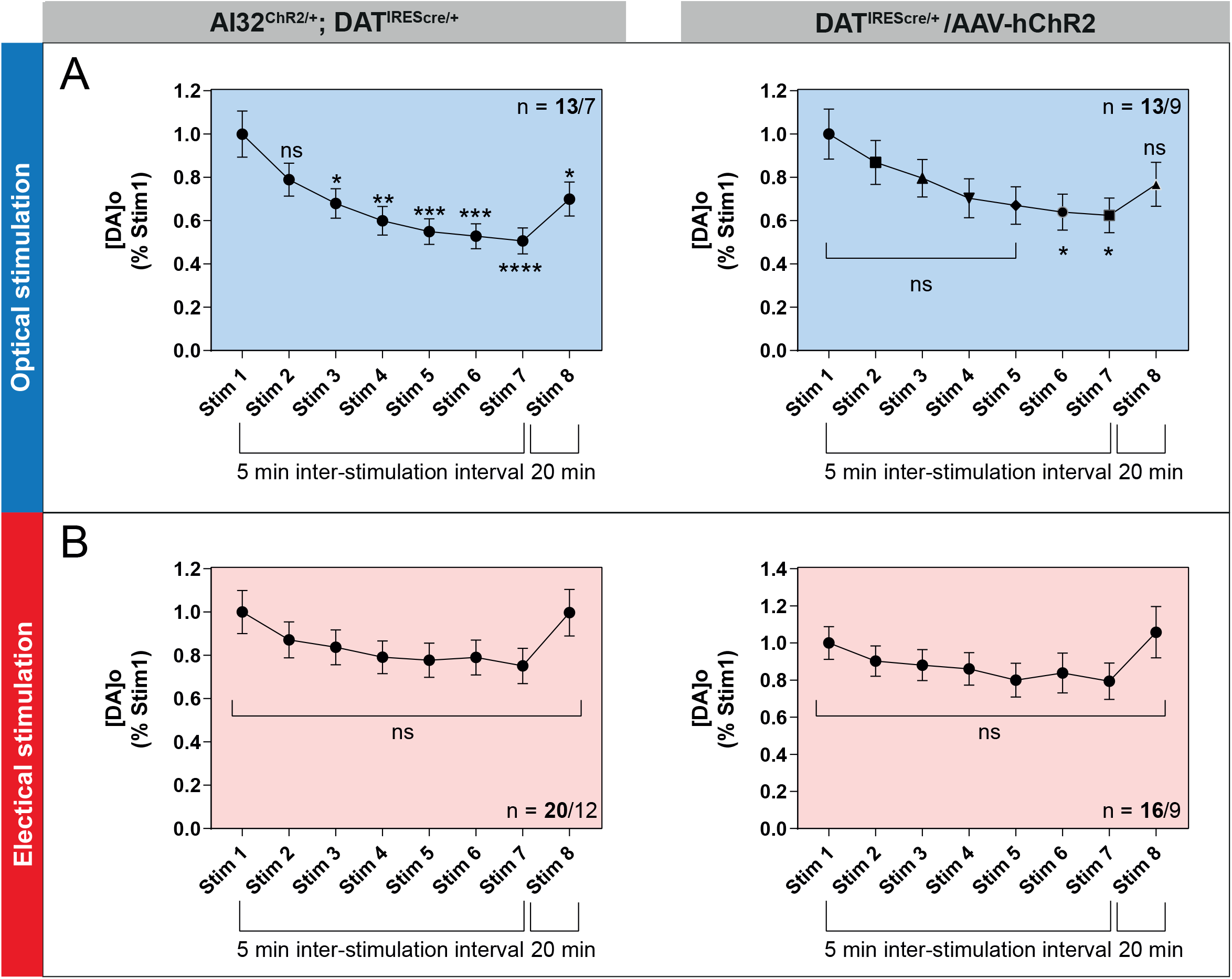
Optogenetic stimulation reveals strong use-dependent attenuation of evoked STD DA release in the VTA. **(A)** Average [DA]o peaks normalized to the first stimulation in the VTA of AI32 and injected DAT^IREScre^ mice evoked by optical stimulation trains (30 pulses of 1 ms at 10 Hz, 470 nm blue light LED). **(B)** Same with pulse-train electrical stimulation (30 pulses of 1 ms at 10 Hz, 400 μA). Each record was obtained in aCSF + 5 μM nomifensine/sulpiride with 1 recording site per slice; inter-stimulus interval between stims 1-7 = 5 min, inter-stimulus interval between stim 7 and 8 = 20 min. Error bars represent +/− S.E.M. and the statistical analysis was carried out by a 1-way ANOVA followed by a Dunnett test (ns, non-significant; *, p < 0.05; **, p < 0.01; ***, p < 0.001; ****, p < 0.0001). The bold number represents the number of slices recorded / number of animals used.

### DA release in the VTA and striatum exhibit similar calcium dependency

As we aimed to examine the role of Syt calcium sensors in STD DA release and in the face of conflicting previous results regarding the extent to which STD DA release depends upon extracellular calcium levels in comparison with axonal release^24,49^, we next evaluated the release of DA at 0, 0.5 and 2.4 mM of extracellular calcium in both the dorsal striatum (axonal release) and VTA (**Fig. 3A**). As expected, based on previous results^13^, no release was detected in the striatum at 0 mM and 0.5 mM calcium, neither in response to single pulses or to trains (n = 13 slices/7 mice) (**Fig. 3B**).

**Figure 3:**
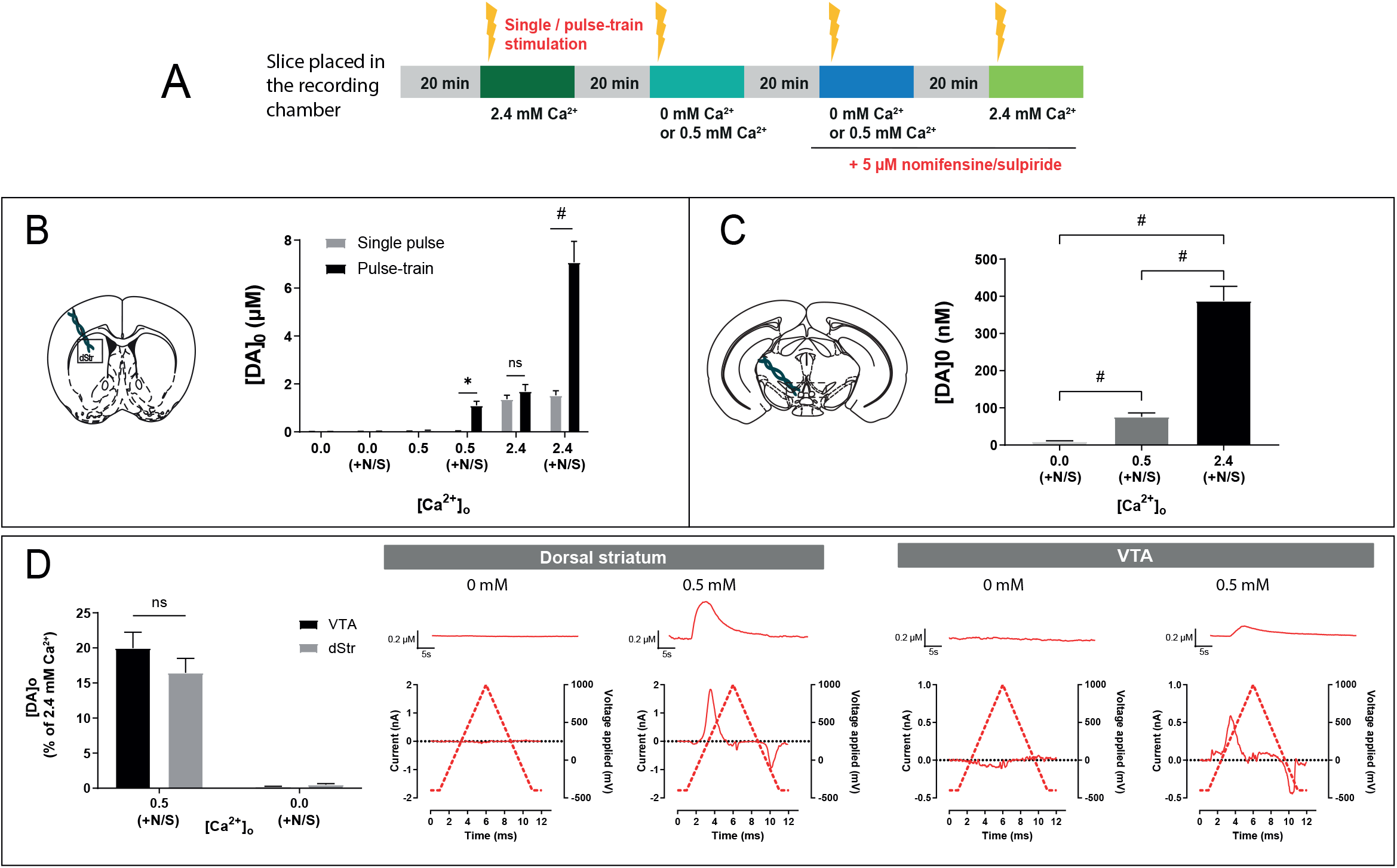
Somatodendritic and axonal dopamine release exhibit a similar calcium dependency. **(A)** Protocol used for FSCV recordings. **(B)** Schematic representation of a striatal slice and average of [DA]o peaks obtained with single or pulse-train stimulations at 0, 0.5 and 2.4 mM of extracellular calcium in the aCSF, with or without addition of 5 μM of nomifensine/sulpiride. **(C)** Schematic representation of a VTA slice and average [DA]o peaks obtained with pulse-train stimulations at 0, 0.5 and 2.4 mM of extracellular calcium in the aCSF containing 5 μM of nomifensine/sulpiride. **(D)** Average [DA]o peaks normalized to 2.4 mM of calcium obtained in the VTA and dorsal striatum (dStr) with pulse-train stimulation and aCSF containing nomifensine/sulpiride. Representative traces and voltammograms are shown on the right. Error bars represent +/− S.E.M. The statistical analysis was carried out by a 1-way ANOVA followed by a Dunnett test (ns, non-significant; ***, p < 0.001; #, p < 0.0001).

In the VTA, STD DA release, here again triggered in the presence of nomifensine/sulpiride (5 μM), was also undetectable at 0 mM extracellular calcium (n = 13 slices/7 animals), but readily detectable at 0.5 mM (n = 16 slices/10 mice) (**Fig. 3C**), as previously described in the guinea pig^49^. Evoked STD DA release at this concentration of calcium was however only 19% of the signal detected at 2.4 mM calcium (76 nM +/−10 nM, compared to 388 nM +/− 39 nM) (**Fig. 3C, 3D**). Recordings performed in striatal slices in the presence of nomifensine and sulpiride similarly revealed detectable DA release at 0.5 mM calcium (**Fig. 3B**) (1.1 μM +/− 0.17 μM, n = 13 slices/7 mice). This represents 16 % of the DA signal detected at 2.4 mM calcium (7 μM +/− 0.87 μM) (**Fig. 3B, 3D**). Therefore, under the same experimental conditions, with no influence of DA uptake and D2 autoreceptor activation, evoked STD and axonal DA release show a similar calcium dependency. All further experiments were performed at 2.4 mM calcium.

### STD DA release is more resilient than axonal release in a Parkinson’s disease model

The differential properties of axonal and STD DA release might in part be involved in explaining the differential impairment of these two forms of release in PD. We therefore examined the resilience of STD DA release in the intrastriatal 6-hydroxydopamine (6-OHDA) model, often used to study adaptation of the DA system in the context of PD progression. Interestingly, previous work performed in the rat showed that extracellular DA levels in the SNc, as determined by *in vivo* microdialysis, are not altered several weeks after a 6-OHDA lesion in the medial forebrain bundle (MFB), while a major loss of DA content was seen in the striatum^33^. We used a protocol adapted from Stott and Barker^34^, who observed that within hours, intra-striatal 6-OHDA (5 μg/μL, 2μL) can impact TH+ fibers in the striatum, while the impact at the STD compartment is delayed for several days post‐surgery, even if approximately 50% of DA neurons degenerate by 12 days. We unilaterally injected saline or 6-OHDA in the dorsal striatum of 6-7-week-old DAT^IREScre^ mice and measured axonal and STD DA release by FSCV 24h or 14 days after the injections (**Fig.4A**).

**Figure 4:**
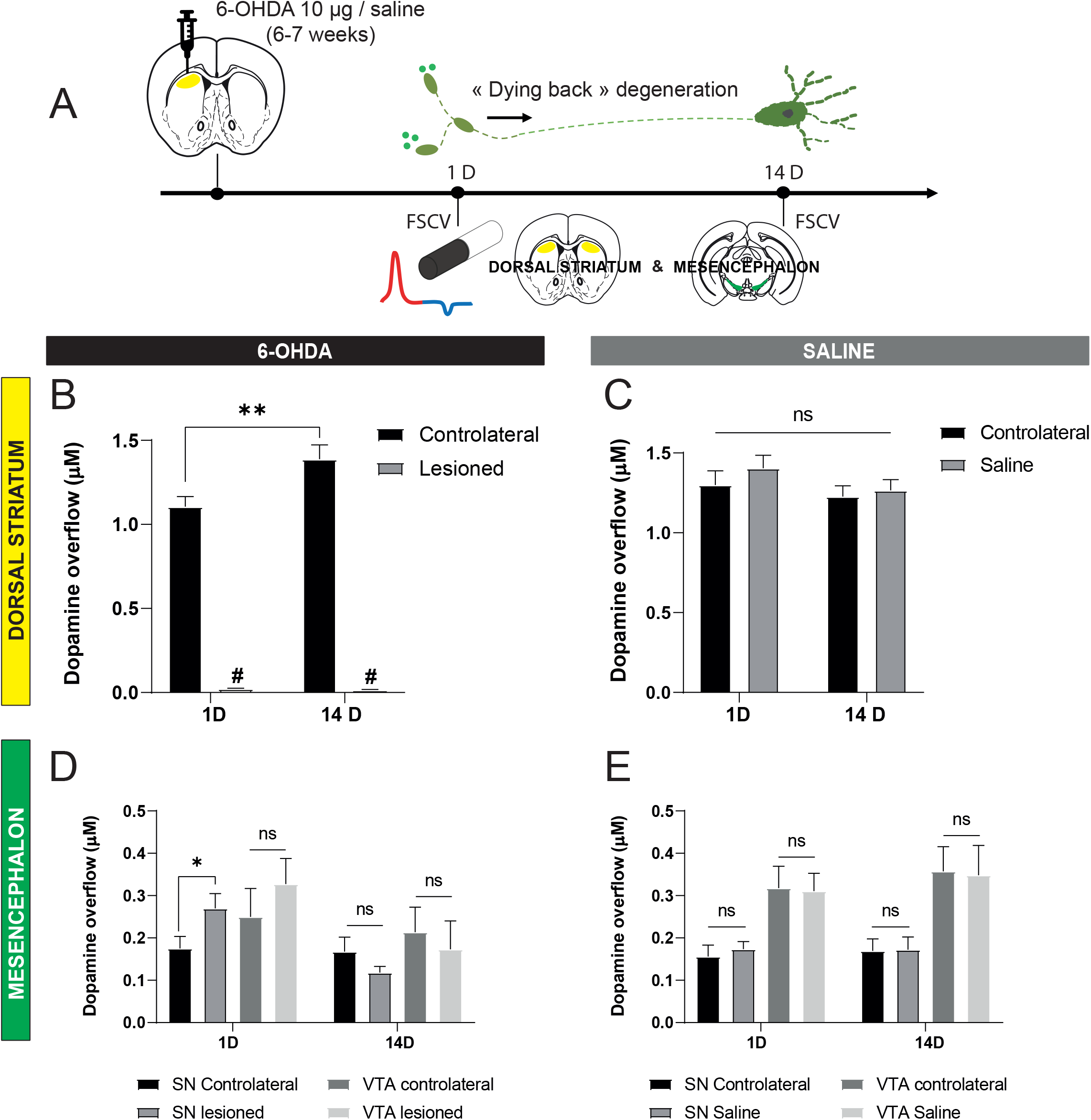
STD DA release is more resilient than axonal release in a model of Parkinson’s disease related axonal dying-back. **(A)** Protocol used for 6-OHDA experiments. Single doses of 2 μl at 5 μg/μl (10 μg) of 6-OHDA were injected in 6-7 weeks old mice in the dorsal striatum. FSCV experiments were conducted 1 day (1D) or 14 days (14D) after the injections and DA overflow was measured in striatal and mesencephalic slices. (**B and C**) Average [DA]o peaks (μM) obtained in the dorsal striatum of 6-OHDA treated (B) and saline-treated (C) mice. (**D and E**) Average [DA]o peaks (μM) obtained in the mesencephalon of 6-OHDA treated (D) and saline-treated (E) mice. Each recording performed in the striatum was obtained in aCSF with an average of 3 recording sites per hemisphere and single pulse stimulation (400 μA, 1 ms). Each recording performed in the SN/VTA was obtained in aCSF + 5 μM nomifensine/sulpiride with pulse-train stimulation (30 pulses, 10 Hz, 400 μA). Error bars represent +/− S.E.M. The statistical analysis was carried out using a t-test (contralateral vs ipsilateral sides) (ns, non-significant; ***, p < 0.001; #, p < 0.0001).

We found at 1-day post-injection a >99% decrease of electrically evoked DA overflow in the dorsal striatum, confirming the robust and acute effect of 6-OHDA on DA axonal fibers [n = 12 slices/6 mice] (**Fig.4B**). This abolition of axonal DA release was also maintained after 14 days [n= 10 slices/5 mice]. Intriguingly, a small decrease of evoked axonal DA overflow was also detected in the contralateral striatum at day-1 (1.1 μM +/− 0.06 μM vs 1.39 μM +/− 0.08 μM at 14 days), something that was not observed in saline-injected control mice (1.3 μM +/− 0.09 μM vs 1.28 μM +/− 0.07 μM at 14 days). As expected, there was otherwise no impact of the injection itself on DA overflow, has seen in all saline treated animals [n = 12 slices/6 mice for each time point] (**Fig.4C and E**).

In a sharp contrast, 1 day after 6-OHDA, STD DA release at the level of the SNc was not reduced, but rather significantly higher in the 6-OHDA lesioned hemisphere (0.27 μM +/− 0.03 μM, [n = 13 slices/6 mice]) compared to the contralateral side (0.18μM +/− 0.03) (**Fig.4D**). At 14 days, a stage at which it is expected that neurodegeneration has reached the STD compartment and approximately half of SNc DA neurons have degenerated^34^, only a tendency of a decrease of DA release in the lesioned-SNc (0.12 μM +/− 0.015, [n = 10 slices/5 mice]) was observed compared to the contralateral side (0.17 μM 0.034 μM), a change that was not significant. There were no significant changes in STD DA release in the VTA region at 1 or 14 days after 6-OHDA. Altogether these data indicate that while axonal release is very sensitive to the toxic effects of a single 6-OHDA injection, STD release is strikingly more resilient.

### Dopamine neurons express the calcium-sensors synaptotagmin 1, 4 and 7

Because Syt1 is the main calcium sensor of axonal release^19,37^, and Syt4 and Syt7 were previously suggested to be critical for STD DA release based on *in vitro* experiments^19^, we next evaluated the presence and subcellular localization of these Syt isoforms in DA neurons *in vivo* in the mouse brain.

Immunohistochemistry was used to test the hypothesis that Syt4 and Syt7 are present within the cell body and dendrites of DA neurons in close association with compartments containing the vesicular monoamine transporter VMAT2. Due to the impossibility to obtain suitable VMAT2 and Syt antibodies produced in different species, we took advantage of a double labelling protocol allowing the use of two primary antibodies from the same species (rabbit) (**Fig. 5A**). The approach was validated by the observation that in control experiments in which the second primary antibody was omitted, no signal was detected for the second antigen, demonstrating that the second secondary antibody was unable to bind to the first primary antibody after the blocking step. Immunoreactivity for Syt4 showed a clear somatic localization in DA neurons, with a notable overlap with VMAT2 (**Fig. 5B**), with little if any signal in terminals in the striatum. Confirming the specificity of the antibody, signal was absent from Syt4 KO DA neurons (**Fig. S2A**). Syt7 immunoreactivity was found in both the STD region of DA neurons as well as in their terminal region in the striatum (**Fig. 5D**). Syt7 immunoreactivity was strongly reduced in Syt7 KO tissue, although some background signal was still detectable (**Fig. S2B**), suggesting sub-optimal specificity. Finally, Syt1 was undetectable in the soma and dendrites of DA neurons, but highly expressed in the terminals in the striatum, as expected (**Fig. 5D**).

**Figure 5:**
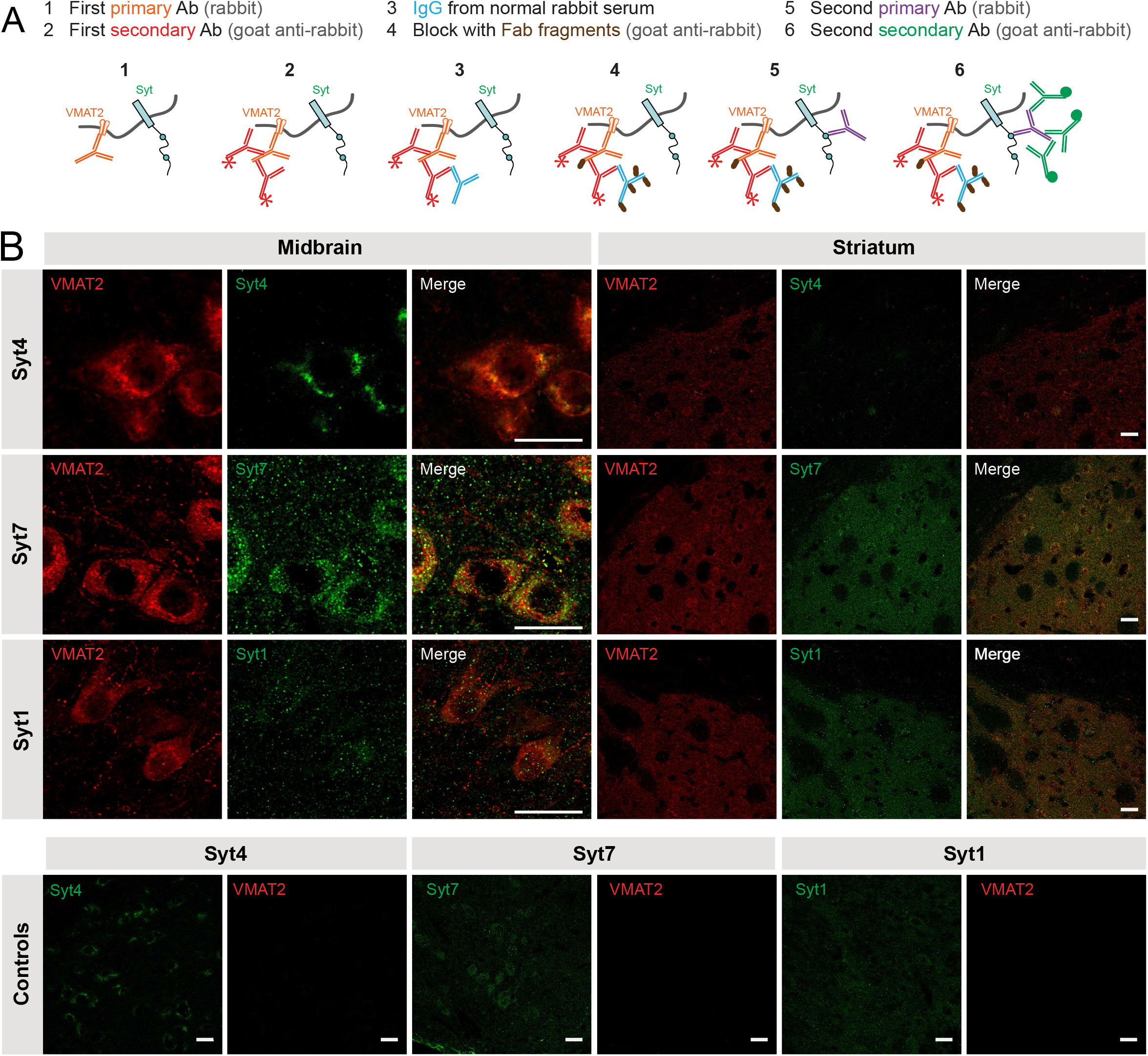
Dopamine neurons express the calcium-sensors synaptotagmin 1, 4 and 7. **(A)** Protocol used for double immunostaining for two primary antibodies from the same host (adapted from a Jackson ImmunoResearch protocol: https://www.jacksonimmuno.com/technical/products/protocols/double-labeling-same-species-primary). Use of normal rabbit serum and unconjugated Fab fragments for blocking after the first secondary. **(B)** Immunohistochemistry of midbrain and striatal slices of adult DAT^IREScre^ heterozygote mice showing colocalization of VMAT2 and either Syt1, Syt4 or Syt7 in DAergic neurons. Scale bar = 20 μm. Control images were obtained using the full protocol without the use of the second primary antibody (in the midbrain).

### Double knockout of Syt4 and Syt7 strongly reduces STD DA release

Considering the expression of Syt4 and Syt7 in DA neurons and their apparent localization in the STD compartment of these neurons, we hypothesized that evoked STD DA release should be reduced in constitutive Syt4 or Syt7 KO mice. These experiments were performed using electrical train stimulation, in the presence of nomifensine and sulpiride. To obtain a thorough understanding of the individual roles of Syt4 and Syt7, wild-type, heterozygous and KO littermates were compared, and recordings were performed for each mouse in the dorsal striatum, the ventral striatum (nucleus accumbens core and shell) and the VTA. These experiments revealed that axonal and STD DA release in the VTA were not significantly reduced in Syt4 or Syt7 KO mice (**Fig. 6A, 6B**). The absence of effect of Syt4 or Syt7 KO on STD DA release could be due to the ability of one isoform to compensate for the other in the STD compartment, with or without compensatory upregulation of the expression of the other isoform. Using qRT PCR in whole brain homogenates, we found that the total levels of Syt7 mRNA in Syt4 KO mice were unchanged, as were the levels of Syt4 mRNA in Syt7 KO mice (**Fig. S2B, S2D**).

**Figure 6:**
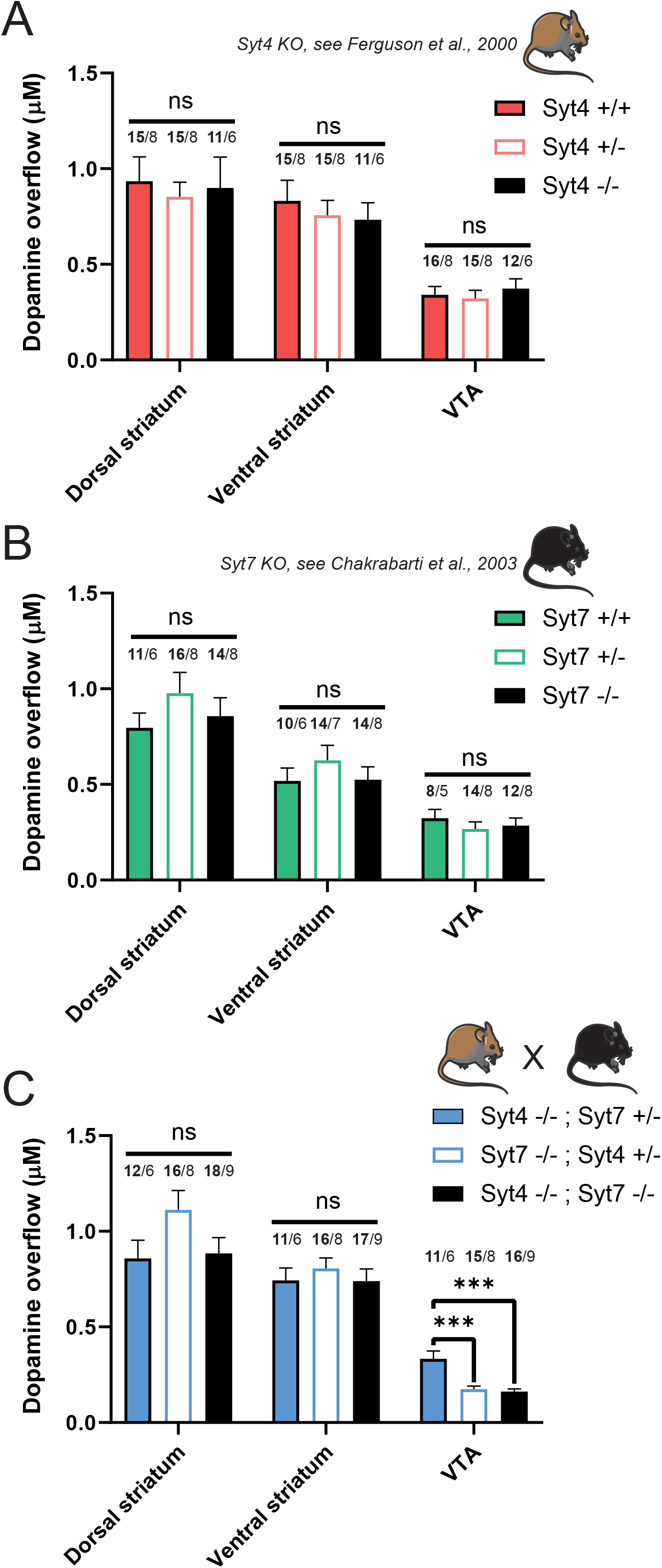
Double KO of Syt4 and Syt7 strongly reduces STD DA release. **(A)** Average [DA]o peaks (μM) obtained in the dorsal striatum, ventral striatum and VTA of Syt4 constitutive knock-out mice bred from heterozygous crosses. **(B)** Same for the constitutive Syt7 knock-out mice. (**C**) Same for double Syt4/Syt7 KO mice (Syt4^-/-^; Syt7^-/-^) and heterozygotes control animals (Syt4^-/-^; Syt7^+/−^ and Syt7^-/-^; Syt4^+/−^). Each recording from the VTA was obtained in aCSF + 5 μM nomifensine/sulpiride with 2 recording sites per slice and pulse-train stimulation (30 pulses, 10 Hz, 400 μA). Error bars represent +/− S.E.M. The statistical analysis was carried out by 1-way ANOVA followed by a Tukey test (ns, non-significant; ***, p < 0.001; # p < 0.0001). The bold number represents the number of slices recorded / number of animals used.

To examine if functional compensation can explain the lack of change in STD DA release in the single KO mice, we next crossed these two mouse lines to generate a double Syt4 and Syt7 KO. As controls, Syt4^-/-^; Syt7^+/−^ and Syt7^-/-^; Syt4^+/−^ animals were also used for the FSCV recordings. Once again, no significant differences were found in the dorsal and ventral striatum (**Fig. 6C**). The amount of DA released in the VTA of Syt4^-/-^; Syt7^+/−^ (0.334 μM +/− 0.04 μM; n = 6 mice) was similar to controls from the Syt4 and Syt7 individual KO mouse lines (respectively 0.341 μM +/− 0.044 μM; n = 8 mice and 0.32 μM +/− 0.046 μM; n = 5 mice). However, in Syt4^-/-^; Syt7^-/-^ animals, we found a robust and significant ≈50% decrease of STD DA release (0.162 μM +/− 0.014 μM; n = 9 mice). Interestingly, the Syt7^-/-^; Syt4^+/−^ animals also showed a systematic decrease of STD DA release of about ~50% (0.175 μM +/− 0.016 μM; n = 8 mice). Together these results argue that both Syt4 and Syt7 isoforms contribute to STD DA release, with functional compensation of one isoform by the other. These data also suggest a more critical role of Syt7 compared to Syt4 because the presence of only one Syt7 allele is sufficient to support STD DA release in the absence of Syt4.

## Discussion

### Characteristics of STD DA release

In the present study, we performed the first characterization of optically evoked STD DA release in the mouse mesencephalon using a combination of optogenetics and FSCV and compared its characteristics to release evoked by electrical stimulation. As previously reported by others, we found that the absolute levels of evoked DA overflow detected in this region were low compared to levels detected in the terminal region in the striatum. Furthermore, we found that a robust STD DA release signal could only be detected using pulse-train stimulation (**Fig.1**). Blocking DA reuptake and D2 autoreceptor function using nomifensine and sulpiride caused a 5-fold increase in peak signal amplitude, thus making detection of this signal straightforward and reproducible.

Using a repeated stimulation protocol, we found that repeated optical stimulation with a 5 min interval produces a strong rundown of STD DA release, whereas no such attenuation was seen with electrical stimulation (**Fig.2**). This finding is compatible with previous results reporting a similar rundown of optically-evoked axonal DA release in the striatum^45^. It is therefore conceivable that the decrease observed with optical stimulation results from a low reserve capacity of STD DA release due to limited vesicular reserve pools in the soma and dendrites of DA neurons. It is equally possible that in response to electrical stimulation, a similar decrement is not observed because this form of stimulation recruits neuromodulatory mechanisms that result from activation of afferent terminals releasing 5-HT, NE, acetylcholine, glutamate, GABA or neuropeptides onto DA neurons^45,50^. Further experiments will be required to test this hypothesis.

Our experiments comparing the impact of changes in extracellular calcium levels of STD and axonal DA release argue for the existence of a similar calcium dependency for both forms of release (**Fig. 3**). These findings are compatible with previous results obtained in mice, which were performed using patch-clamp recordings and the measurement of STD D2 receptor^24^. It is possible that a different conclusion was reached in guinea pig brain slices because some aspect of the STD DA release mechanism is different in that species^13^. Another possibility is that a small component of axonal DA release is also included in the signal detected in the VTA. This possibility has been raised previously^49^, but the available anatomical data actually suggests that DA containing axonal varicosities are extremely scarce in the VTA^51,52^, except in the context of compensatory axonal sprouting associated with partial lesions^53^. It would nonetheless be useful to revisit this question with additional anatomical work in the future to provide more quantitative data.

### Implications of STD DA release in PD

The differential resilience of STD DA release in comparison to that of terminal DA release is of particular interest because a major hypothesis of PD progression proposes that loss of function begins at the axon terminal level, only later progressing to loss of cell bodies (i.e. the dying back hypothesis of PD)^32,54^. In this context, it may be hypothesised that at early stages of PD, STD DA release may still be functional and contribute to partial maintenance of DA-dependent regulatory mechanisms in the ventral midbrain. Of relevance, it has been proposed that STD DA release contributes, along with axonal DA release to motor behaviors^55,56,10,57^. Although a major focus of PD research has been on restoring DA release in the striatum with L-DOPA treatment^58^ or with transplantation of mesencephalic tissue^59^, the possible contribution of STD DA release to functional adaptation or perturbation of basal ganglia circuit function in PD has received little attention until now. Here we thus evaluated how STD DA release changes over time after intra-striatal 6-OHDA, used to model PD axonal dying-back. We observed that axonal release is very sensitive to the neurotoxic effect of 6-OHDA, as previously known. In comparison, we found that STD DA release persists with no major decrement for up to 14 days after the lesion, thus reflecting its high level of resilience. This observation is in agreement with previous data from microdialysis experiments measuring basal DA levels in the mesencephalon of 6-OHDA-lesioned rats^33^. Our finding of an increase in STD DA release in the SNc at 1 day after the lesion further suggests that at early stages of PD pathophysiology, the loss of axonal DA signaling in the dorsal striatum could constitute a signal for SNc neurons to upregulate their STD DA release as a possible compensatory mechanism to sustain normal functions. Although this hypothesis is speculative and will require further experiments to clarify the full time-course and the mechanisms involved, these findings raise interest in further investigating STD DA release and its plasticity in more disease-relevant PD models. It would also be of interest to disentangle the possible contributions of STD and axonal DA release in the VTA after partial lesions because of the possible appearance of aberrant compensatory axonal fibers in the mesencephalon in such models^53^.

### Contribution of Syt4 and Syt7 to STD DA release

Finally, we examined the contribution of the synaptotagmin isoforms Syt4 and Syt7 to STD DA release. Acute downregulation of both isoforms has previously been shown *in vitro* to severely reduce STD DA release, with no similar effect of Syt1 downregulation^19^. Although our present immunostaining results provide further support for the presence of these proteins in the STD compartment of DA neurons, we failed to detect any significant decrease in evoked STD DA release in VTA slices prepared from individual constitutive Syt4 or Syt7 KO mice. It is possible that contrarily to acute downregulation with siRNAs, constitutive gene deletion may lead to homeostatic compensation leading to elevated levels of Syt4 in Syt7 KO mice and vice versa. It is also possible that *in vitro* models lack homeostatic compensatory mechanisms that are recruited *in vivo*. Our experiments quantifying Syt4 and Syt7 total mRNA levels failed to provide support for such a compensation. Another possibility is therefore that Syt4 and Syt7 play similar roles in supporting STD DA release and that one can compensate for absence of the other in the context of constitutive gene deletion. The robust decrease in activity-dependent STD DA release in Syt4/Syt7 double KO mice supports this interpretation. In Syt4^-/-^; Syt7^-/-^ mice or Syt7^-/-^; Syt4^+/−^ mice, we observed a two-fold decrease of STD DA release. This decrease was surprisingly not found in Syt4^-/-^; Syt7^+/−^ mice, strongly suggesting that Syt7 plays a particularly important role and that a single allele of Syt7 is sufficient to sustain STD DA release in the absence of Syt4.

Considering that in the absence of both Syt4 and Syt7, approximately half of total STD DA release levels remain, we hypothesise that other calcium sensors are also involved. One possible candidate is Syt11. This isoform is interesting because like Syt4, it has been reported to be present in the STD compartment of neurons^60^. Like Syt4, it also contains a natural mutation in one of its C2 calcium-binding domain, compatible with a regulatory role in exocytosis rather than a classical calcium-sensing role^61^. Finally, like Syt4, Syt 11 is a risk locus for PD^62,63^, and is a substrate of the E3 ubiquitin ligase parkin, linked to early-onset familial forms of PD. Intriguingly, Syt11 overexpression in the SNc has been reported to cause a decrease of DA release in the striatum^64^. Finally, the main synaptotagmin isoform Syt1 might also be of interest, as it was recently demonstrated as the main calcium sensor for fast striatal DA release^37^. We have not found strong evidence for localization of Syt1 in the STD domain of DA neurons, but further examination of this possibility with higher resolution techniques would be warranted. A broader evaluation of the contribution of other synaptic and exocytosis proteins in STD DA release would also be useful. Interestingly, evoked STD DA release measured as D2-IPSCs was recently reported to be abolished in mice with conditional deletion of the active zone protein RIM, while spontaneous release remained intact^65^. However, the subcellular localization of RIM in the STD compartment of DA neurons is currently undetermined.

Together our work provides a new perspective on the mechanisms of STD DA release and renews the interest in better understanding its roles in normal brain function and in diseases such as PD.

---

**Supplementary figure 1:**
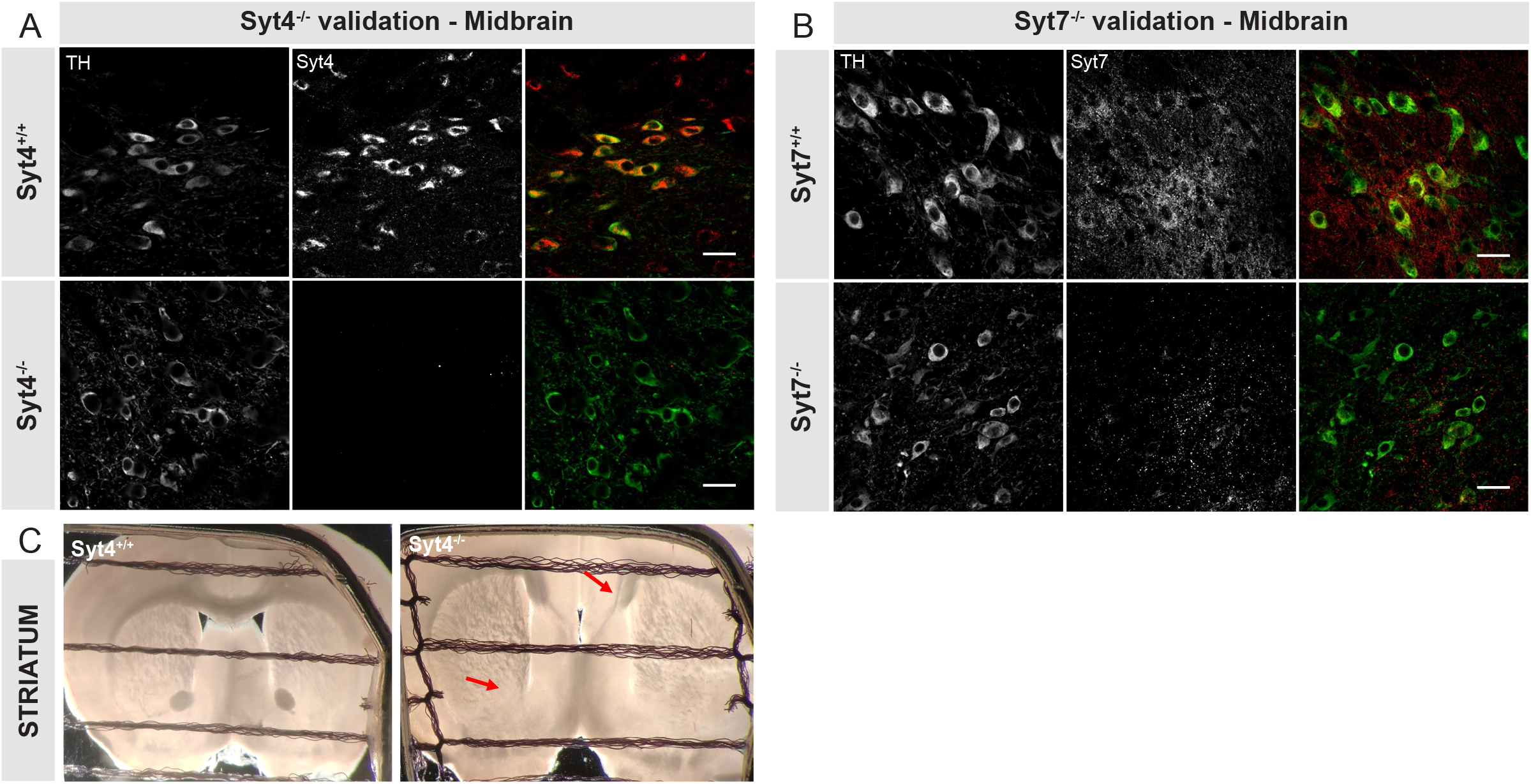
Knockout validation of Syt4 and Syt7 antibodies. **(A)** TH and Syt4 immunostaining of adult Syt4 WT (+/+) and KO (-/-) mesencephalon showing the specificity of the Syt4 antibody. (**B)** TH and Syt7 immunostaining of adult Syt7 WT (+/+) and KO (-/-) mice showing a strong reduction in signal, with some remaining background signal in the KO animals. The anti-Syt7 antibody was generated against a recombinant peptide comprising amino acids 46–133 of the unique Syt7 spacer domain ^66^. The targeting vector generated a stop codon after the position coding for amino acid 83 in exon 4 (**Fig. S2**), thus making it possible that the remaining signal corresponds to a short, mutated protein comprising the lumenal domain, the transmembrane region, and only a fraction of the spacer domain. (**C**) Representative striatal brain slices recorded during FSCV experiments in WT and Syt4 constitutive KO mice. Red arrow indicates obvious neurodevelopmental defects at the level of the anterior commissure and the corpus callosum in KO animals.

**Supplementary figure 2:**
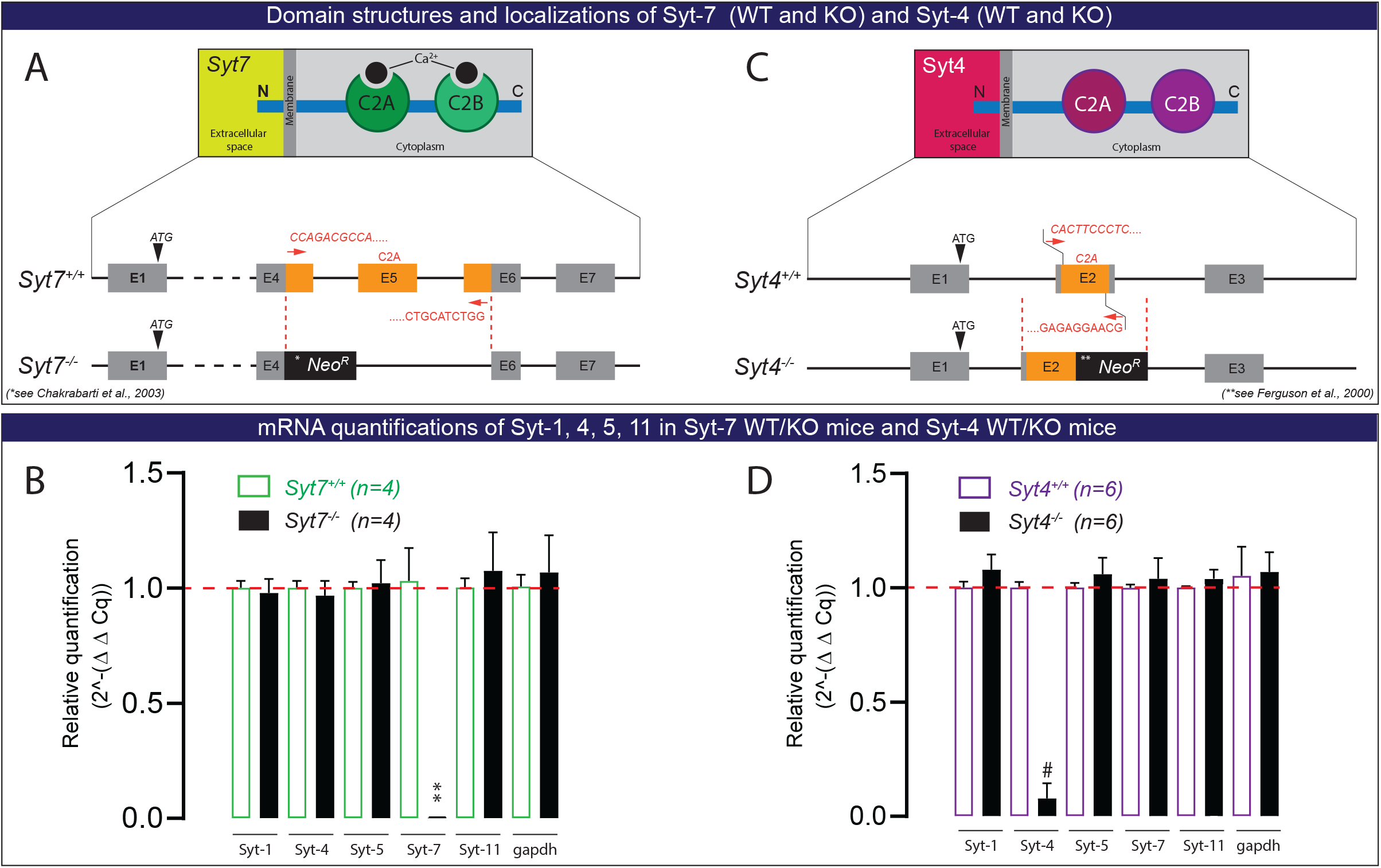
No compensatory changes in Syt1, Syt4, Syt7, Syt11 mRNA in Syt4 and Syt7 constitutive KO mice. (**A and B**) Schematic representation of the construction of Syt4 and Syt7 KO (-/-) mice. The primers used in qRT PCR for amplifying the deleted region of each gene are indicated. (**C and D**) Relative changes of mRNA levels measured by qRT-PCR in Syt7 (C) and Syt4 (D) KO mice. Ct values (mean of duplicate repeats) of Syt1, 4, 5, 7 and 11 mRNA levels were normalized to the Ct value of GAPDH in the same samples. Error bars represent +/− S.E.M. The statistical analysis was carried out using a t-test (WT vs KO samples) (**, p < 0.01; #, p < 0.0001).

## Author contributions

All authors participated in the design of experiments, data analysis, and interpretation. CD performed qRT PCR experiments. NF performed double immunostaining experiments. WK designed genotyping primers. BDL and LET wrote the manuscript.

## Acknowledgements

We thank Dr. Wade G Regehr (Harvard Medical School) for providing the Syt7 constitutive KO mice, Dr. Gary Miller (Columbia University) for providing the VMAT2 antibody and Marie-Josée Bourque for managing the mouse colonies and performing mouse genotyping. This work was funded by the National Sciences and Engineering Research Council of Canada (NSERC, grant RGPIN-2020-05279) to LET. LET also received support from the Krembil Foundation, the Brain Canada Foundation and the Henry and Berenice Kaufmann Foundation. BDL received a graduate student award from Parkinson Canada. CD received a studentship from the Fonds de la Recherche du Québec en Santé (FRQS).

